# FAP20 is required for flagellum assembly in *Trypanosoma brucei*

**DOI:** 10.1101/2024.01.19.576295

**Authors:** Michelle M. Shimogawa, Keya Jonnalagadda, Kent L. Hill

## Abstract

*Trypanosoma brucei* is a human and animal pathogen that depends on flagellar motility for transmission and infection. The trypanosome flagellum is built around a canonical “9+2” axoneme, containing nine doublet microtubules (DMTs) surrounding two singlet microtubules. Each DMT contains a 13-protofilament A-tubule and a 10-protofilament B-tubule, connected to the A-tubule by a conserved, non-tubulin inner junction (IJ) filament made up of alternating PACRG and FAP20 subunits. Here we investigate FAP20 in procyclic form *T. brucei*. A FAP20-NeonGreen fusion protein localized to the axoneme as expected. Surprisingly, FAP20 knockdown led to a catastrophic failure in flagellum assembly and concomitant lethal cell division defect. This differs from other organisms, where FAP20 is required for normal flagellum motility, but generally dispensable for flagellum assembly and viability. Transmission electron microscopy demonstrates failed flagellum assembly in FAP20 mutants is associated with a range of DMT defects and defective assembly of the paraflagellar rod, a lineage-specific flagellum filament that attaches to DMT 4-7 in trypanosomes. Our studies reveal a lineage-specific requirement for FAP20 in trypanosomes, offering insight into adaptations for flagellum stability and motility in these parasites and highlighting pathogen versus host differences that might be considered for therapeutic intervention in trypanosome diseases.

**SIGNIFICANCE STATEMENT:**

- Diverse eukaryotic organisms rely on a generally conserved axoneme architecture and dynein-dependent beating mechanism to drive motility, but mechanisms conferring lineage-specific motility needs are largely unknown.
- FAP20 is a conserved flagellar protein that impacts flagellum motility in multiple organisms.
- The current work demonstrates FAP20 is particularly important in the pathogen, *T. brucei*, providing insight into pathogen adaptations for moving through host environments and illuminating targets to consider for therapeutic intervention in trypanosome diseases.

## INTRODUCTION

*Trypanosoma brucei* and related African trypanosomes are deadly pathogens of humans and other mammals and have historically presented a tremendous burden on human health and prosperity throughout sub-Saharan Africa (Kristjanson et al., 1999, Abro et al., 2021). While dedicated efforts and development of new therapeutic options (Alvarez-Rodriguez et al., 2022, Valverde Mordt et al., 2022) have reduced the human health burden, these parasites remain a threat to the lives of 70 million people living in 36 countries (Papagni et al., 2023). Related kinetoplastid parasites, *Trypanosoma cruzi* and *Leishmania* spp., afflict almost a billion people worldwide (Rao et al., 2019). Meanwhile, *T. brucei* infection of agriculturally important livestock presents a substantial economic burden, with an estimated annual unrealized income of almost five billion dollars in endemic areas (Abro et al., 2021). Animals also serve as reservoirs of infection that limit efforts to eradicate the disease (Kasozi et al., 2023). Beyond their direct medical and economic importance, trypanosomatid parasites are valuable models for understanding fundamental eukaryote cell biology (Serricchio and Bütikofer, 2011, Vincensini et al., 2011, Lukeš et al., 2023). As such, there is great interest in understanding trypanosome biology.

*T. brucei* is a flagellated protozoan that is transmitted between mammalian hosts by a tsetse fly vector. Flagellum-mediated motility is required for transmission through the tsetse and for infection of a mammalian host (Rotureau et al., 2014, Shimogawa et al., 2018). *T. brucei* is an exclusively extracellular parasite and flagellated at all developmental stages throughout its life cycle. Its flagellum must be stable enough to support vigorous movement while laterally attached to the cell body, and withstand external forces encountered while moving through blood and tissues of the mammalian host and insect vector (Langousis and Hill, 2014, Imhof et al., 2019, Zhang et al., 2021). Beyond its role in motility, the *T. brucei* flagellum also directs cell morphogenesis (Moreira-Leite et al., 2001) and provides a platform for assembling signaling machinery that mediates host-pathogen interactions necessary for transmission and pathogenesis (Oberholzer et al., 2011, Salmon et al., 2012, Shaw et al., 2019, Velez-Ramirez et al., 2021, Bachmaier et al., 2023). Therefore, understanding flagellum structure and composition in trypanosomes has become a topic of great interest (Sáez Conde and Dean, 2022).

The trypanosome flagellum is composed of a canonical “9+2” axoneme, together with a lineage-specific paraflagellar rod (PFR) filament that is laterally connected to the axoneme along most of its length (Langousis and Hill, 2014). The 9+2 axoneme consists of 9 outer doublet microtubules (DMTs) arranged around 2 singlet microtubules (Fig. 1A). Each microtubule is made from α/β-tubulin dimers that polymerize end-to-end to form protofilaments, which in turn connect laterally to form a cylinder (Downing and Nogales, 1998). DMTs contain one complete, 13-protofilament A-microtubule (A-tubule), connected to an incomplete, 10-protofilament B-microtubule (B-tubule) ((Nicastro et al., 2011), Fig. 1A). Recent work in trypanosomes and other organisms has provided insight into proteins that mediate DMT assembly and function (Stoddard et al., 2018, Ichikawa et al., 2019, Imhof et al., 2019, Ma et al., 2019, Owa et al., 2019, Gui et al., 2021, Gui et al., 2022, Li et al., 2022, Kubo et al., 2023, Shimogawa et al., 2023). Of particular interest is the inner junction (IJ) that connects the B-tubule to the A-tubule at the inner face of each DMT (Yanagisawa et al., 2014, Dymek et al., 2019, Khalifa et al., 2020, Shimogawa et al., 2023). The IJ contains a non-tubulin filament together with microtubule inner proteins (MIPs) that interconnect with the IJ filament and each other, facilitating attachment of protofilament 10 in the B-tubule to protofilament 1 of the A-tubule (Yanagisawa et al., 2014, Dymek et al., 2019, Khalifa et al., 2020, Shimogawa et al., 2023) (Fig. 1A). Unlike protofilaments, the IJ filament is a non-tubulin polymer of alternating PACRG and FAP20 subunits (Yanagisawa et al., 2014, Dymek et al., 2019). Most organisms contain a single PACRG, while some, such as trypanosomes and *Tetrahymena* (Kubo et al., 2023), contain two PACRG genes (Dawe et al., 2005, Kubo et al., 2023).

**Figure 1.**
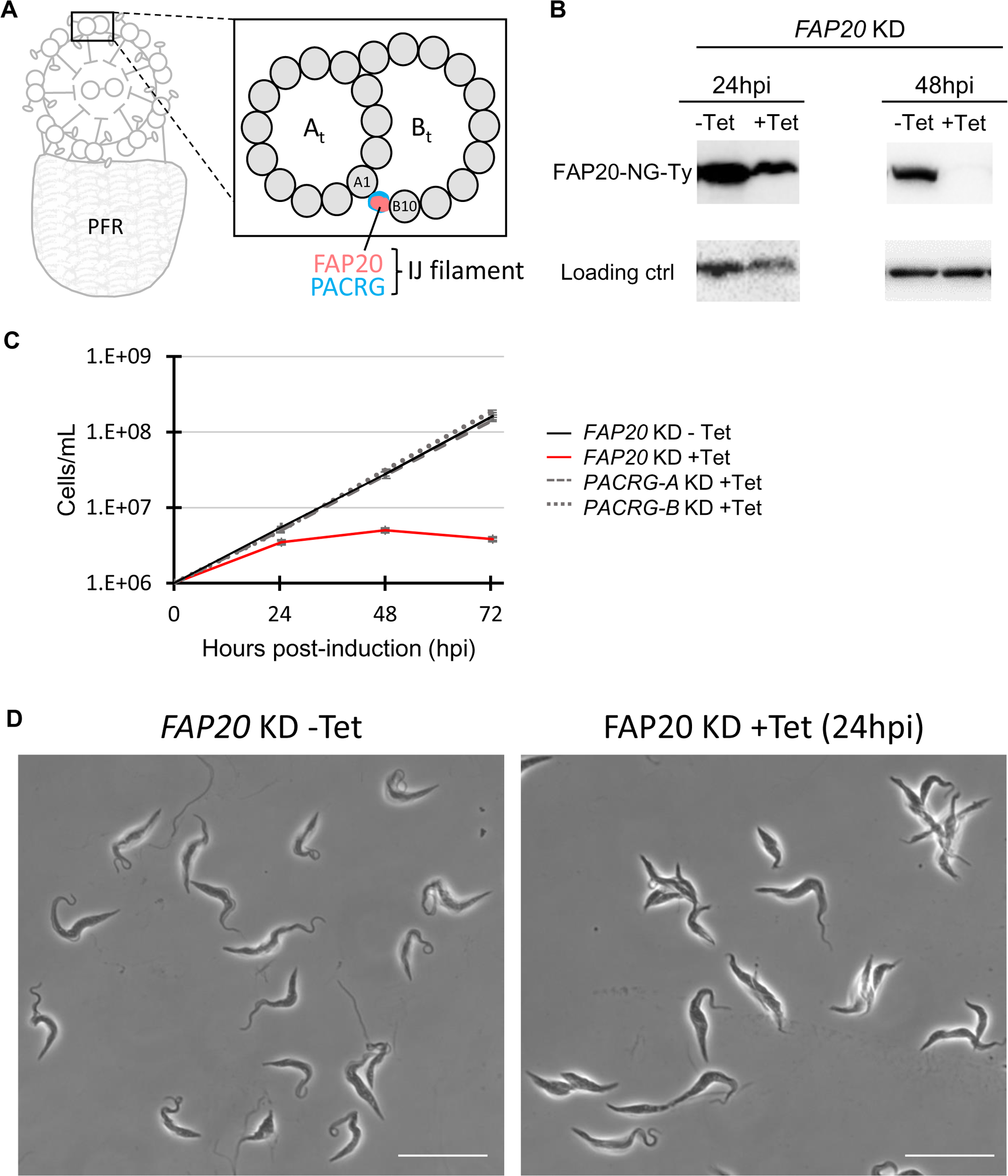
*FAP20* KD is lethal in *T. brucei*. (*A*) Cartoon showing a cross-section of the *T. brucei* flagellum comprised of a 9+2 axoneme and paraflagellar rod (PFR). Inset shows a cross-section of a doublet microtubule (DMT) with the A- and B-microtubules (At, Bt) labeled and tubulin protofilaments in grey. The inner junction (IJ) filament formed by FAP20 and PACRG connects tubulin protofilaments A1 and B10. (*B*) *FAP20* knockdown (*FAP20* KD) parasites were grown in the presence of tetracycline (Tet) to induce knockdown. Western blot analysis of FAP20 protein expression, as detected with anti-Ty antibody, or anti-tubulin (24hpi) or anti-PFR2 (48hpi) as loading controls. 5×10^6^ cell equivalents were loaded per lane for anti-Ty (24hpi and 48hpi) and the 48hpi loading control (PFR2). 2.5×10^6^ cell equivalents per lane were loaded for the 24hpi loading control (tubulin). (*C*) Cumulative growth curve shows the mean cell density ± standard deviation vs time from two independent biological replicates (*FAP20* KD) or three independent biological replicates (*PACRG-A* KD and *PACRG-B* KD). (*D*) Microscopy of fixed *FAP20* KD cells grown with or without Tet. Scale bar = 20 μm.

Studies in several organisms indicate PACRG and FAP20 are required for IJ filament assembly and axoneme motility, but generally dispensable for overall flagellum assembly (Laligne et al., 2010, Mendes Maia et al., 2014, Yanagisawa et al., 2014, Dymek et al., 2019, Chrystal et al., 2022, Gonzalez-Del Pozo et al., 2022, Chen et al., 2023). In *Chlamydomonas* for example, the DMT can even assemble with overall normal architecture in the complete absence of the IJ filament (Dymek et al., 2019). It is likely therefore, that MIP proteins around the IJ cooperate with the IJ filament to maintain B-tubule architecture and attachment to the A-tubule. In *T. brucei*, knockdown of either of the two PACRG homologues, PACRG-A or -B, did not affect cell growth or DMT assembly, while knockdown of both together slowed cell growth and led to DMT assembly defects that became progressively worse toward the distal end of the axoneme, although cells remained viable and overall flagellum length was not affected (Dawe et al., 2005). FAP20, the remaining IJ filament subunit, has not previously been studied in *T. brucei* and is the focus of the current work.

Here, we investigate *T. brucei* FAP20 function using a combination of inducible RNAi knockdown, fluorescence microscopy and thin-section transmission electron microscopy (TEM). Our results show that FAP20 is essential for flagellum assembly and cell viability, suggesting the unique importance of FAP20 and the IJ filament for flagellum stability and motility in this parasite.

## RESULTS AND DISCUSSION

### *FAP20* knockdown is lethal in *T. brucei*

To study FAP20 function, we used tetracycline (Tet)-inducible RNAi knockdown (KD) to generate a ‘*FAP20* KD’ cell line. RNAi was performed in procyclic cultured form (PCF) cells expressing an mNeonGreen (NG)-tagged version of FAP20 (FAP20-NG) and knockdown was confirmed by Western blot (Fig. 1B). FAP20 protein was reduced within 24 hours post-induction (hpi) and not detected by Western at 48hpi (Fig. 1B). To our surprise, *FAP20* knockdown was rapidly lethal, with cell doubling ceasing between 24 and 48hpi (Fig. 1C; Movie 1). An independent cell line, targeting the *FAP20* 3’UTR (‘*FAP20* UTR KD’), exhibited the same lethal phenotype (Movie 2). This result contrasts with what is observed following knockdown of either of the other two IJ subunits, PACRG-A or -B, which had no effect on cell growth (Fig. 1C, Supp. Fig. 1), as reported previously (Dawe et al., 2005). Prior studies showed that simultaneous knockdown of both *PACRG-A* and *-B* did slow cell growth, although the defect was not evident until approximately 72hpi and cells remained viable (Dawe et al., 2005). The rapid, lethal phenotype of *FAP20* KD also differs from knockdown of *T. brucei* MIPs examined to date, which have minimal impact on parasite growth or viability (Shimogawa et al., 2023).

To better understand the nature of the lethal phenotype, we examined *FAP20* KD and *FAP20* UTR KD cells directly. Microscopic examination of intact cells revealed that *FAP20* knockdown resulted in clumps of two or more cells that failed to separate, in some cases with abnormal morphologies (Fig. 1D). Cell motility was severely compromised upon knockdown (Movies 1 and 2). Control cells had vigorously beating flagella and generally moved propulsively with the flagellum tip leading, but *FAP20* KD and *FAP20* UTR KD cells were essentially immotile (Movies 1 and 2). On occasion, one cell in a clump retained an actively beating flagellum, but the others generally showed no indication of flagellar motility at all (Movies 1 and 2). This is one of the most severe motility defects described for knockdown of flagellar proteins in *T. brucei*, as most mutants lack normal cell propulsion but retain a beating flagellum (Bastin et al., 1998, Branche et al., 2006, Baron et al., 2007, Baron et al., 2007, Absalon et al., 2008, Rotureau et al., 2014).

The cell division phenotype in *FAP20* knockdowns bears some resemblance to what has been reported previously for other flagellar mutants in PCF cells (Branche et al., 2006, Ralston et al., 2006, Baron et al., 2007). Those mutants typically start cytokinesis and then fail at the final stage of cell septation, yielding clusters of multiple cells having mostly normal morphology, but connected at their posterior ends. *FAP20* KD cells, in contrast, fail at an earlier stage of cleavage furrow ingression and cells in clumps often remain laterally connected with abnormal morphology, reminiscent to what is reported for bloodstream form (BSF) flagellar mutants (Ralston and Hill, 2006). The combined defects – lethality, failure in cytokinesis, and nearly immotile flagella – demonstrate a critical requirement for FAP20 in flagellum function in this organism.

### *FAP20* KD blocks flagellum assembly in *T. brucei*

*T. brucei*’s prominent flagellum is clearly visible as a dark line in phase contrast images of detergent-extracted cytoskeletons (Fig. 2) and *FAP20*-NG is localized along the flagellum (Fig. 2, (Billington et al., 2023)), as expected. FAP20-NG is lost in most cells of the *FAP20* KD, although some cells retain signal even at 48hpi (Fig. 2), which indicates FAP20 does not readily turn over once incorporated into the flagellum, as described for *Chlamydomonas* FAP20 (Yanagisawa et al., 2014). Interestingly, in cells lacking FAP20-NG, the flagellum appears short or absent (Fig. 2), indicating loss of FAP20 disrupts assembly of the entire flagellum. We therefore used immunofluorescence against the paraflagellar rod (PFR) as a flagellum marker (Saada et al., 2014). Samples were examined at 24hpi because that is the first timepoint assessed where the growth defect was evident. In control cells, FAP20-NG signal localized alongside the PFR, starting near the distal tip of the flagellum and extending further than the PFR at the flagellum’s proximal end, as expected for an axonemal protein. In *FAP20* KD cultures, cells without FAP20-NG have a short and sometimes misshapen PFR (Fig. 3), demonstrating that flagellum assembly is ultimately blocked. In cells that retained FAP20-NG, the FAP20-NG signal stopped substantially before the end of the PFR at the distal tip of the flagellum (Fig. 3, arrowheads), presumably corresponding to cells in which PFR assembly continued after *FAP20* knockdown. These data further distinguish the *FAP20* KD PCF phenotype from that of *PACRG-A/B* double mutants (Dawe et al., 2005), and typical flagellar mutants (Baron et al., 2007, Shimogawa et al., 2023), as these generally assemble a normal length PFR and axoneme (Dawe et al., 2005, Baron et al., 2007, Shimogawa et al., 2023). Moreover, the data also distinguish the PCF *FAP20* KD phenotype from that of BSF flagellar mutants, which fail at cleavage furrow ingression but, in contrast to *FAP20* KD, continue through multiple rounds of axoneme and PFR replication (Ralston and Hill, 2006). The combined data demonstrate that FAP20 is critical for stable assembly of the entire flagellum and that short segments of the PFR can assemble without FAP20, but this is not sustained.

**Figure 2.**
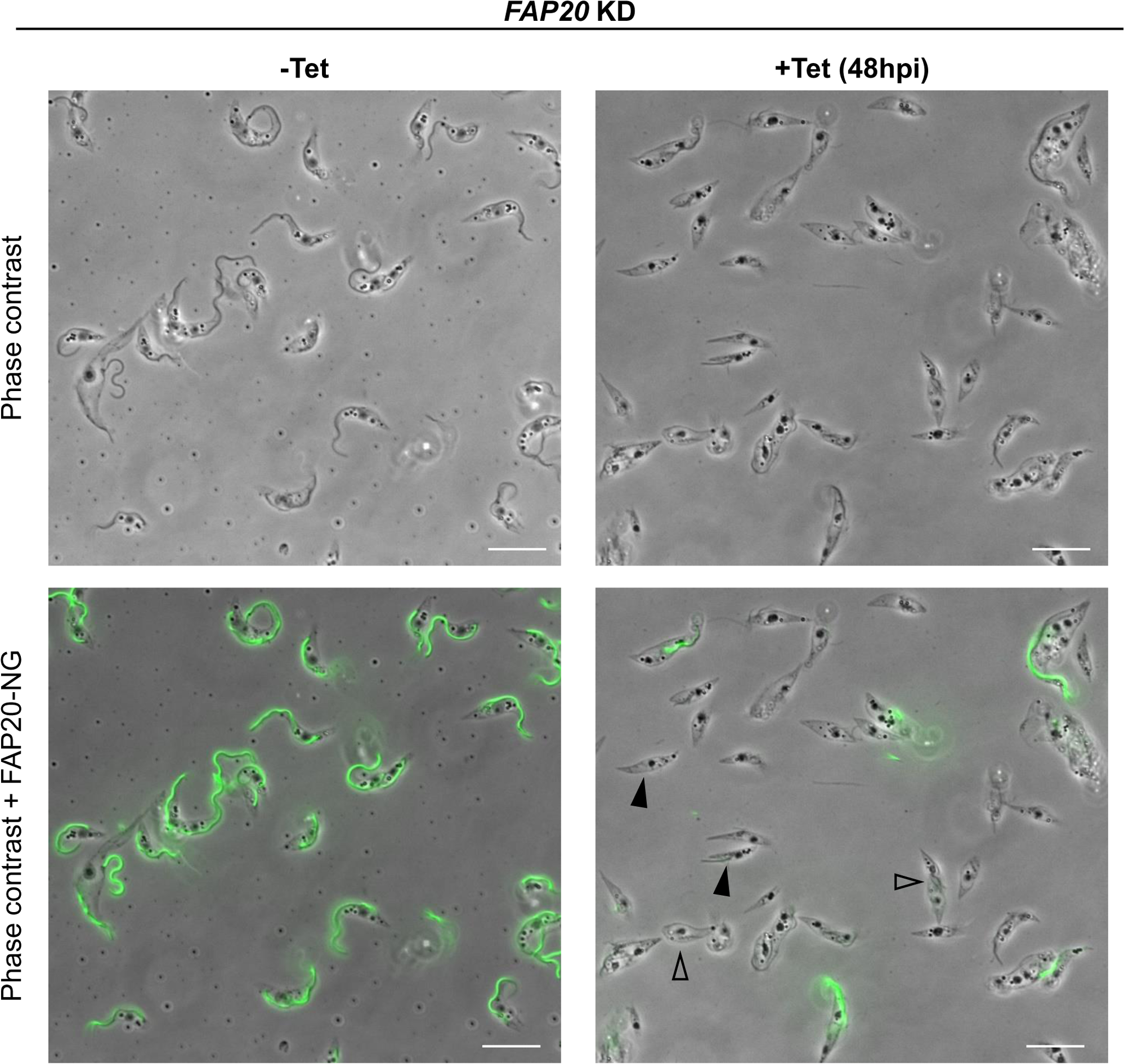
*FAP20* KD causes severe morphological defects. *FAP20* was knocked down by tetracycline (Tet)-inducible RNAi in cells expressing FAP20-NG (green). Detergent-extracted cytoskeletons were prepared from uninduced *FAP20* KD cells (-Tet) or cells grown 48h in Tet to induce KD (+Tet). Open arrowheads show cells that are enlarged or failed in cell division. Black arrowheads show cells with short or absent flagella.

**Figure 3.**
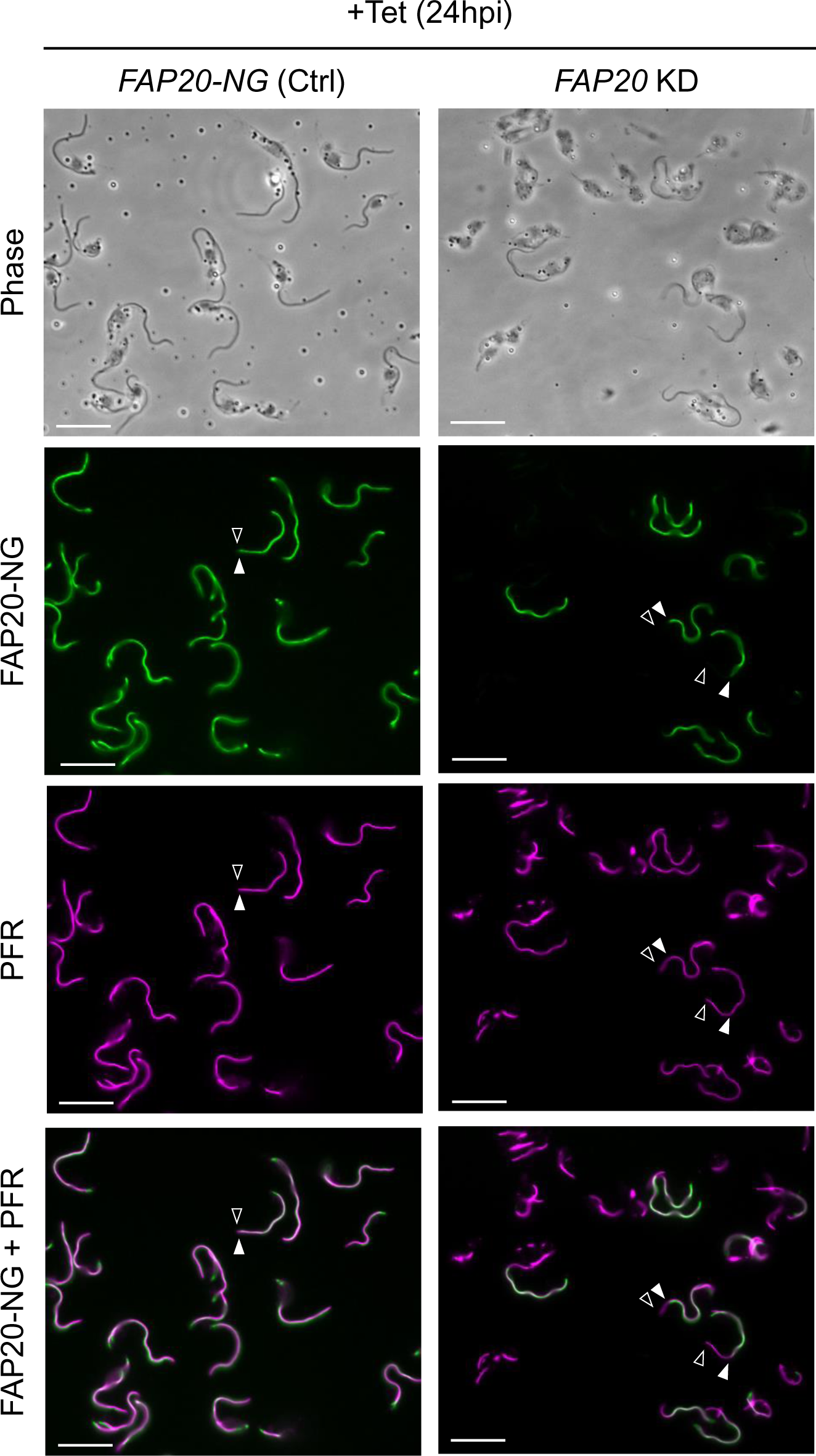
*FAP20* KD disrupts flagellum assembly. Detergent-extracted cytoskeletons were prepared from FAP20-NG (Ctrl) and *FAP20* knockdown (*FAP20* KD) parasites grown in tetracycline (Tet) for 24h to induce knockdown. Images show mNeonGreen (NG)-tagged FAP20 (green) and paraflagellar rod (PFR, magenta). The distal end of FAP20 signal is indicated by a closed arrowhead, while the distal end of PFR signal is indicated with an open arrowhead. Scale bar = 20 μm. Images are representative of at least two independent experiments.

### FAP20 is required for stable flagellum assembly uniquely in *T. brucei*, indicating a critical role for the IJ in this pathogen

To elucidate ultrastructural foundations of the *FAP20* KD flagellum assembly defect, we examined cells by thin section transmission electron microscopy (TEM). At low magnification, the flagellum defect is clearly evident, as fewer cell bodies had associated flagella in *FAP20* KD versus control sections (Fig. 4A; 15/25 = 60% for *FAP20* KD vs 29/34 = 85% for ctrl). When examined at higher resolution, control cells were seen to contain a canonical 9 + 2 arrangement of axonemal microtubules (Fig. 4B). *FAP20* KD cells, on the other hand, exhibited a range of doublet microtubule defects, including singlets, incomplete doublets, displaced doublets, and extranumerary doublets (Fig. 4B). Occasionally we observed PFR defects, e.g. incomplete or excessive PFR material (Fig. 4B), consistent with PFR immunofluorescence results (Fig. 3). Axonemal defects were primarily restricted to doublet microtubules and obvious defects in the central pair were rarely observed, except in sections where the entire axoneme was distorted.

**Figure 4.**
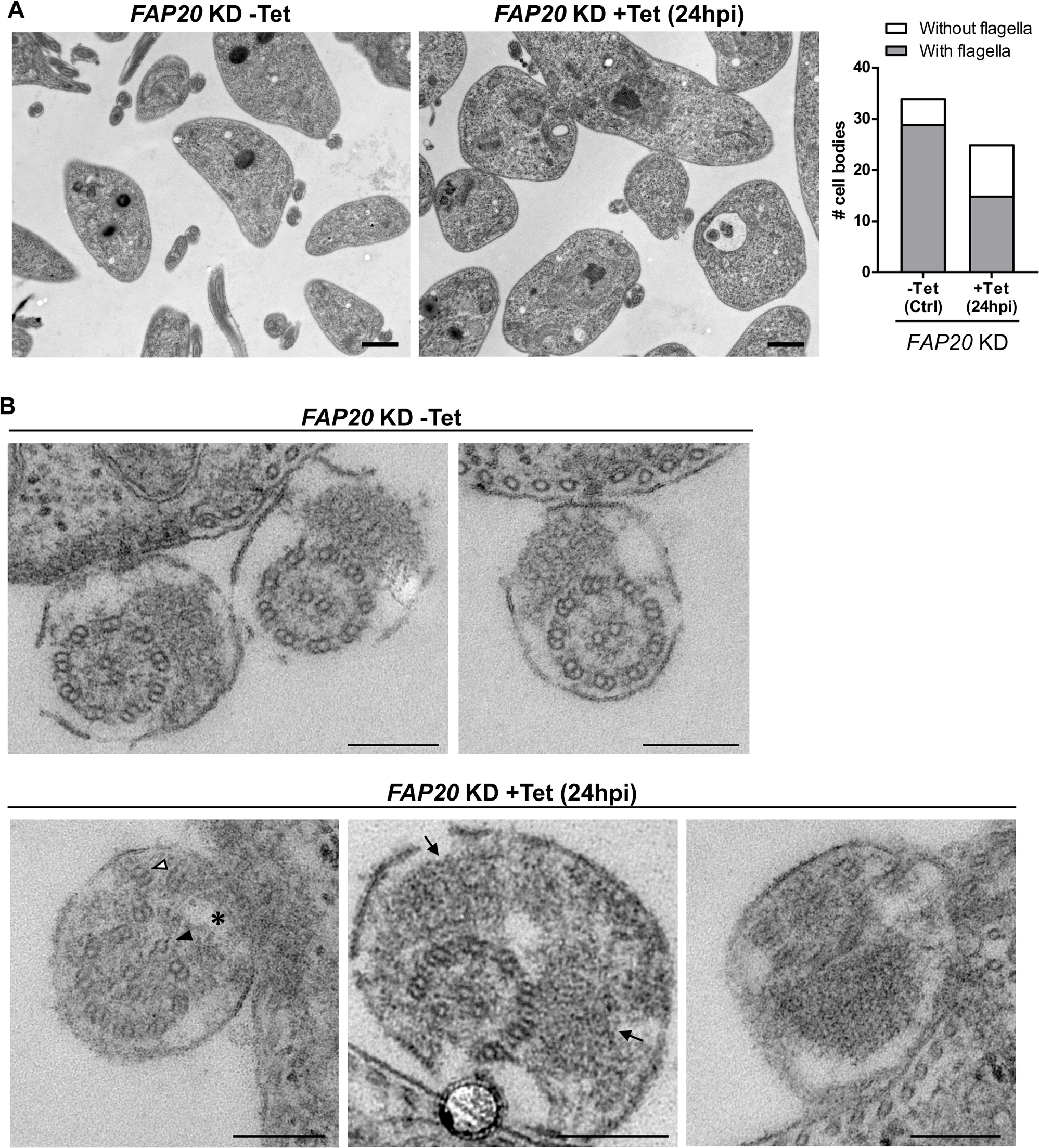
*FAP20* KD disrupts flagellum ultrastructure. (*A*) Representative thin section transmission electron microscopy (TEM) images of *FAP20* KD cells without tetracycline (-Tet) and after 24h Tet-induction. Scale bar = 1 µm. Images are representative of two independent experiments. The number of cell bodies with an associated flagellum was quantified (10/25 = 40% without flagella for *FAP20* KD +Tet versus 5/34 = 15% without flagella for *FAP20* KD -Tet). (*B*) TEM images of individual flagella, showing examples of displaced, extranumerary doublets (open arrowhead), singlets or incomplete doublets (arrowheads), incomplete PFR (*) or excessive PFR (arrows). Scale bar = 200 nm. Images are representative of two independent experiments.

Our combined data demonstrate that loss of FAP20 is lethal in *T. brucei* due to failure of flagellum assembly. Cryogenic electron microscopy (CryoEM) analysis indicates FAP20 of *Chlamydomonas* is found at both the IJ of the DMT and at the base of the C2a projection of the central pair complex (CPC) (Yanagisawa et al., 2014, Dymek et al., 2019, Gui et al., 2022). While we cannot formally distinguish between a CPC defect versus an IJ defect as the source of the *FAP20* KD phenotype, TEM data demonstrate prominent DMT defects and little to no detectable disruption of the CPC (Fig. 4B). Moreover, knockdown of CPC proteins in *T. brucei* does not block flagellum assembly and is not lethal (Branche et al., 2006, Ralston et al., 2006). Therefore, we favor the idea that flagellum assembly failure and concomitant lethal phenotype are due to requirement of FAP20 at the IJ.

FAP20 function has been studied in several organisms, where mutants show defective ciliary motility, perturbation of the IJ filament, and reduced axoneme stability, albeit without blocking assembly of the flagellum, which grows to mostly normal length and in some cases longer (Laligne et al., 2010, Mendes Maia et al., 2014, Yanagisawa et al., 2014, Dymek et al., 2019, Chrystal et al., 2022, Gonzalez-Del Pozo et al., 2022, Chen et al., 2023). Recent studies have shown that FAP20 can directly connect tubulin dimers to intact microtubules *in vitro*, and enhance microtubule stability (Bangera et al., 2023). These *in vitro* studies provide a biochemical basis for FAP20 in connecting tubulin protofilaments of the A- and B-tubules at the IJ and promoting axoneme stability. Notably, however, in no case in other organisms did loss of FAP20 block flagellum assembly. In many instances the DMT exhibited mostly normal architecture despite absence of FAP20 and, in some cases, even complete absence of the entire IJ filament (Dymek et al., 2019). Thus, in other organisms, FAP20 contributes to but is not required for DMT assembly and stability, whereas loss of FAP20 in *T. brucei* leads to rapid and complete failure of stable flagellum assembly and is lethal. Indeed, this is one of the most severe defects reported for a flagellar protein in PCF *T. brucei* cells (Dawe et al., 2005, Branche et al., 2006, Baron et al., 2007, Ralston and Hill, 2008). A potential explanation for lineage-specific dependency on FAP20 in *T. brucei* is the lineage-specific poly-glutamine (poly-Q) C-terminal extension that is uniquely found on the *T. brucei* protein compared to FAP20 from all other organisms in which FAP20 function has been studied (Fig. 5). Given the extensive interconnections among IJ subunits and MIPs of the DMT (Khalifa et al., 2020, Shimogawa et al., 2023), together with numerous lineage-specific trypanosome MIPs (Imhof et al., 2019, Shimogawa et al., 2023), it is conceivable that loss of FAP20 disrupts a trypanosome-specific interaction with other trypanosome proteins critical for flagellum assembly. As has been noted previously (Heddergott et al., 2012, Bargul et al., 2016, Imhof et al., 2019) several constraints, e.g. attachment to the PFR and cell body, vigorous, three-dimensional beating, and encounters with host tissues, impose unique demands on axoneme stability in trypanosomes (Imhof et al., 2019). These constraints might make the need for FAP20 in supporting axoneme stability particularly acute in trypanosomes.

**Figure 5.**
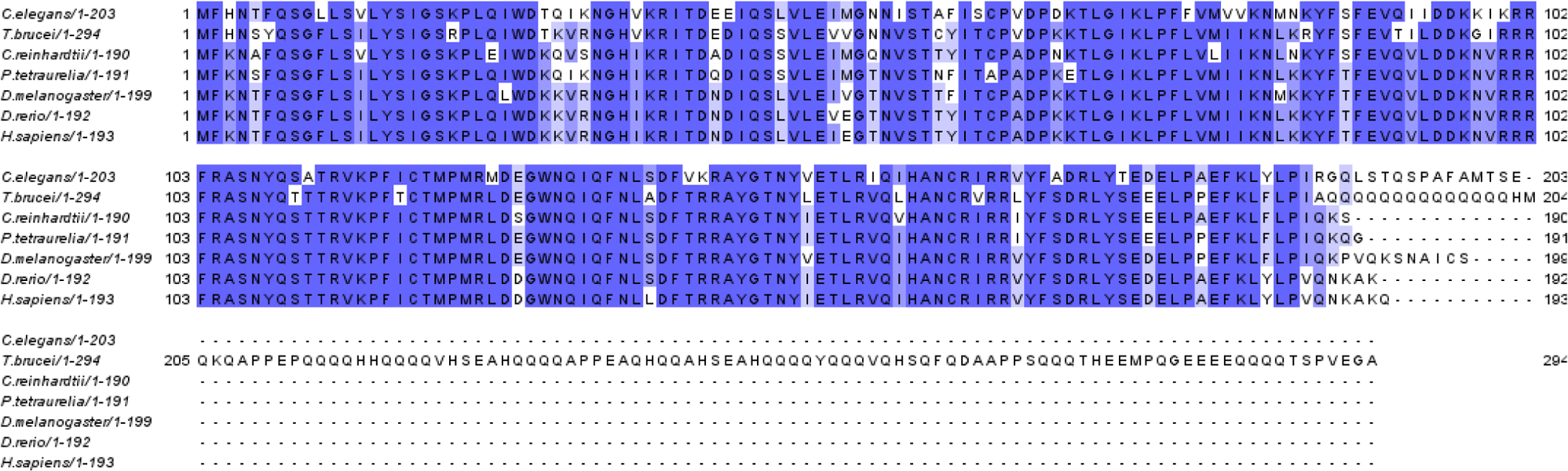
Clustal Omega alignment of FAP20 protein sequences in which FAP20 function has been studied. *Trypanosoma brucei* FAP20 has a long C-terminal poly-glutamine (poly-Q)-rich domain that is not present in organisms that do not require FAP20 for flagellum assembly.

While the underlying basis for FAP20-dependency in trypanosomes remains to be determined, our combined results underscore the importance of FAP20, and the IJ filament, for flagellum stability and motility in trypanosomes relative to other flagellated eukaryotic cells, and suggest FAP20 may accommodate unique adaptations required for the parasite to move through its insect vector and vertebrate host. As trypanosomes are medically and economically important pathogens, our results highlight a difference between pathogen and host that might be exploited for therapeutic intervention.

## MATERIALS AND METHODS

### Biological materials

All novel biological materials are available from the authors upon request.

### Trypanosoma brucei culture

Procyclic *T. brucei brucei* (strain 29-13) originally obtained from George Cross (Rockefeller University) (Wirtz et al., 1999) were cultivated in SM medium (Oberholzer et al., 2009) supplemented with 10% heat-inactivated fetal bovine serum (FBS) at 28°C with 5% CO2.

### In situ tagging

mNeonGreen (NG)-tagged cell lines were generated in the 29-13 background by C-terminal tagging (Dean et al., 2015) with pPOTv6-puro-puro-NG. See Supp. Table 1 for list of primers.

### Tetracycline-inducible knockdown

Constructs for tetracycline-inducible knockdown were designed using RNAit (Redmond et al., 2003) and cloned into p2T7-177 (Wickstead et al., 2002). See Supp. Table 1 for list of primers. NotI-linearized plasmids were transfected into the corresponding NG-tagged cell lines using established methods (Oberholzer et al., 2009). Clonal lines were generated by limiting dilution. Knockdown was induced by growing cells in the presence of 1 µg/mL tetracycline (Tet). For growth curves, cells were counted with a Beckman Coulter Z1 particle counter and diluted daily to a concentration of 1×10^6^ cells/mL.

### Western blotting

Cells were washed in Dulbecco’s phosphate buffered saline (DPBS) and boiled in 1x Laemmli sample buffer (Bio-rad). Samples were separated on a 10% acrylamide gel and transferred to nitrocellulose membrane. Membranes were blocked in phosphate buffered saline (PBS) + 5% milk and incubated overnight at 4°C in primary antibody diluted in blocking solution (1:3000 mouse anti-Ty BB2 (Bastin et al., 1996), 1:10,000 mouse anti-beta tubulin E7 (Developmental Studies Hybridoma Bank), or 1:10,000 rabbit anti-PFR2 (Saada et al., 2014)). Membranes were washed three times in PBS + 0.05% Tween-20 (PBS-T) and incubated in goat anti-mouse or goat anti-rabbit horseradish peroxidase (HRP)-conjugated secondary antibody (Bio-rad) diluted 1:5000 in blocking solution. After washing three times in PBS-T, membranes were developed using the SuperSignal West Pico PLUS chemiluminescent substrate kit (Thermo) and images were captured on a Bio-rad ChemiDoc MP imaging system.

### Video microscopy

Cells were applied to a glass slide and covered with a glass coverslip. *FAP20* KD videos were acquired at ∼30fps on a Zeiss Axiovert 200M microscope with a 40x objective lens using a Basler ace acA5472-17um camera and Pylon software. Videos were cropped in Fiji and played back at 30fps. *FAP20* UTR KD time lapse images were acquired at ∼1fps on a Zeiss Axioskop II microscope with a 63x objective lens using a Zeiss Axiocam 705 camera and Zen software. Images were cropped in Zen and converted into 30fps videos in Fiji.

### Microscopy of fixed whole cells

Cells were washed in PEME buffer (100 mM PIPES, 2 mM EGTA, 1 mM MgSO4, 0.1 mM EDTA, pH 6.8) and resuspended in DPBS. Cells were fixed with 0.2% paraformaldehyde on ice for 5 min, washed in DPBS and allowed to dry onto poly-L-lysine treated coverslips. Coverslips were incubated in −20°C methanol for 30s, dried, and rehydrated in DPBS before mounting on slides. Images were acquired on a Zeiss Axiskop II microscope with a Plan-Apochromat 63x/1.4 objective lens using Zen software.

### Fluorescence microscopy of detergent-extracted cytoskeletons

Cells were washed in PEME buffer, resuspended in DPBS and allowed to settle onto poly-L-lysine treated coverslips for 10 min. Non-adhered cells were removed and cytoskeletons were extracted on the coverslip for 5 min in PEME + 1% IGEPAL CA-630 (NP-40). Coverslips were rinsed in PEME and mounted directly on microscope slides or processed for immunofluorescence to detect the paraflagellar rod (PFR), as follows. Coverslips were blocked in DPBS + 8% normal donkey serum + 2% bovine serum albumin for 45 min, then incubated with 1:1000 rabbit anti-PFR2 primary antibody (Saada et al., 2014) diluted in blocking solution for 1.5 hr. Coverslips were washed three times for 10 min in DPBS + 0.05% Tween-20 (DPBS-T) and incubated with 1:1500 donkey anti-rabbit Alexa 594 secondary antibody (Invitrogen) diluted in blocking solution for 1 hr. Coverslips were washed three times as above, rinsed in DPBS and mounted on slides. Images were acquired on a Zeiss Axiskop II microscope with a Plan-Apochromat 40x/1.4 or 63x/1.4 objective lens or a Zeiss Axio Imager Z1 fluorescence microscope with a Plan-Apochromat 63x/1.4 objective lens using Zen software.

### Thin section transmission electron microscopy (TEM)

Cells were fixed directly in cell culture medium by addition of 2.5% glutaraldehyde (Electron Microscopy Sciences) and pelleted. Cell pellets were resuspended in 4% paraformaldehyde + 2.5% glutaraldehyde in 0.1 M sodium cacodylate buffer, pH 7.2 (Electron Microscopy Sciences) and incubated for 1 hr. Cells were pelleted and resuspended in fresh fixative solution and processed as follows at the Electron Core Facility, UCLA Brain Research Institute (all reagents and materials from Ted Pella). After wash, samples were embedded in 4% agarose gel and post-fixed in 1% osmium tetroxide. After wash, samples were dehydrated through a graded series of ethanol concentrations and propylene oxide. After infiltration with Eponate 12 resin, the samples were embedded in fresh Eponate 12 resin and polymerized at 60°C for 48h. Ultrathin sections of 70nm thickness were prepared and placed on formvar carbon coated copper grids and stained with uranyl acetate and lead citrate. The grids were examined using a JEOL 100CX transmission electron microscope at 60 kV and images were captured by an AMT digital camera (Advanced Microscopy Techniques Corporation, model XR611). Two independent biological replicates were fixed and examined for each sample (−/+ Tet).

### Quantification of flagella in TEM sections

Four to six randomly-acquired images for each sample were captured at low magnification (2900x). The number of cell bodies with or without an associated flagellum was quantified and the data were plotted using Graphpad Prism. Only cell bodies that were completely within the field of view (to allow unambiguous assessment of an associated flagellum) were counted.

## ACKNOWLEDGMENTS

Funding was provided by NIH grant AI052348 (K.L.H.). KJ was supported by the Beckman Scholars Program (Beckman Foundation). We thank Dr. Chunni Zhu at the UCLA Brain Research Institute Electron Microscopy Core Facility for TEM analysis and Angeline Wijono for technical assistance. We thank Elissa Hallem and Peter Bradley for access to their camera and microscope for collecting videos and microscopy images.

