## Supplementary material for "FAP20 is required for flagellum assembly in *Trypanosoma brucei*": Supp Fig 1 and Supp Table 1

Supplementary Figure 1

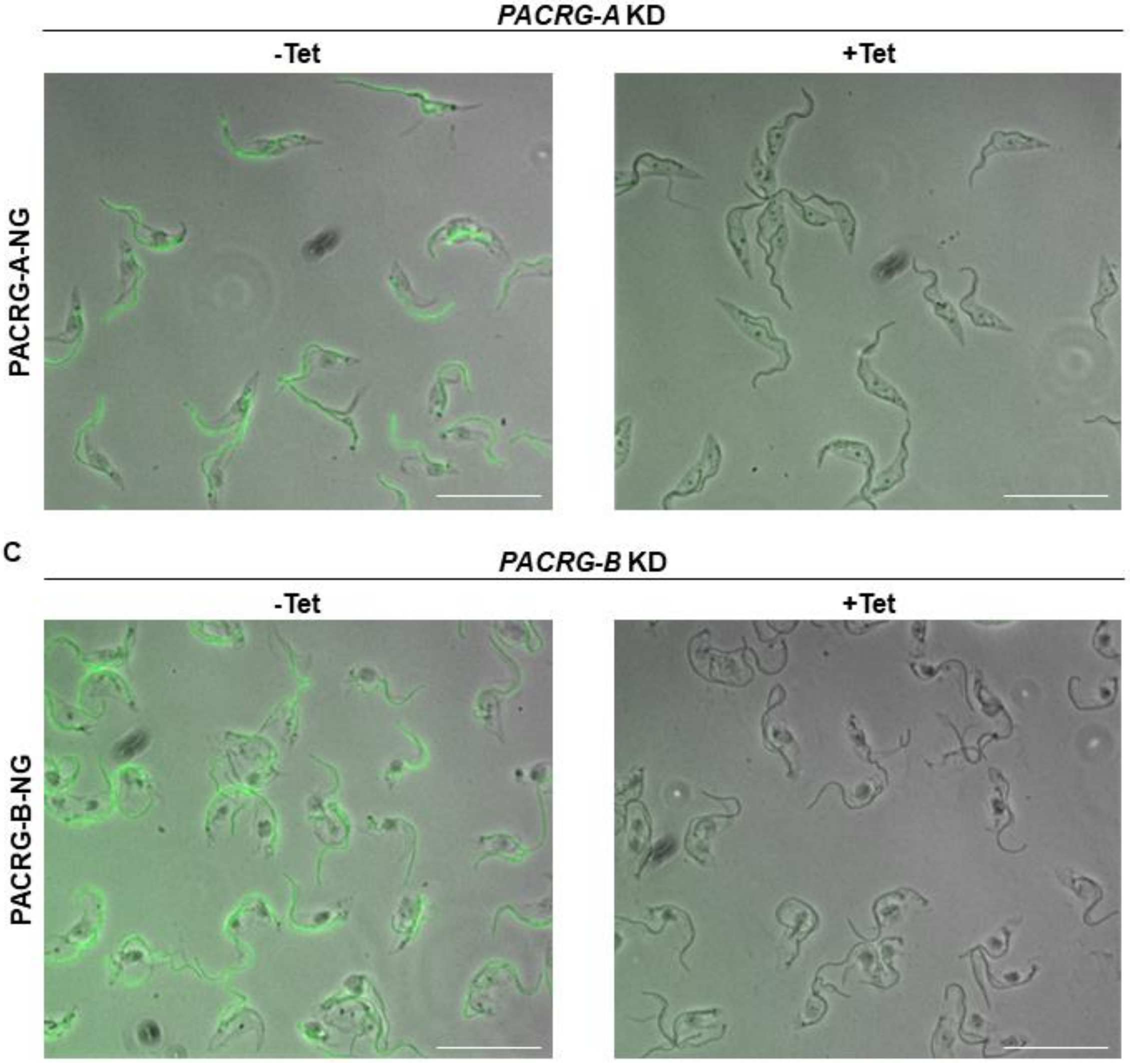


**Supplementary Figure 1. Tetracycline (Tet)-inducible knockdown (KD) of *T. brucei PACRG* homologs.** (*A*) *PACRG-A* and (*B*) *PACRG-B* were independently knocked down by Tet-inducible RNAi in cells expressing the respective target proteins tagged with mNeonGreen (NG). Detergent-extracted cytoskeletons were prepared from uninduced cells (-Tet) or cells in which KD was induced for 72h (+Tet). Scale bars = 20 μm.

**Supplemental Table 1: Primers**

| | | Restriction Site | Primer Sequence |
|---|---|---|---|
| FAP20-NG tagging | Forward | | CCCAACAGCAGACACATGAGGAAAT<br>GCCTCAGGGTGAAGAGGAGGAACAG<br>CAGCAACAAACGTCACCAGTAGAGG<br>GCGCTCTTGTCTCGAAAGGTGAGGA |
| FAP20-NG tagging | Reverse | | ACAGAAACTCGCACGCGAGCGCATA<br>AACACACACACACACAAAAAAAAACA<br>GTCTGCGCACCATTTTTATATGTTCC<br>CCTCCAATTTGAGAGACCTGTGC |
| FAP20 KD | Forward | XbaI | GCtctagaATGACAAGGGCATTCGTCGT |
| FAP20 KD | Reverse | HindIII | CCCaagcttCTTGTTGCTGTTGGTGAGC<br>C |
| FAP20 UTR KD | Forward | XbaI | GCtctagaATGGTGCGCAGACTGTTTTT |
| FAP20 UTR KD | Reverse | HindIII | CCCaagcttGTGCGGGTATGAGAACAT<br>GC |
| PACRG-A-NG tagging | Forward | | TGTTGGAGGTTCATGGTGGCGACGA<br>TGCGTATATCAACATCAAATATATGGT<br>GCCGACGTATGAGAGCTGTATTTTCA<br>GTCTTGTCTCGAAAGGTGAGGA |
| PACRG-A-NG tagging | Reverse | | GGGTGAGCTTTCTTTTCTAACTTCAAA<br>AAACGCATCTTTTCTCATTCTTTCTTC<br>TCGACGTTCCTCCGGCATCACGCTG<br>CCCAATTTGAGAGACCTGTGC |
| PACRG-A KD | Forward | XbaI | GCtctagaGGCACCACGACATCCTTGTA |
| PACRG-A KD | Reverse | HindIII | CCCaagcttCGCGCTTCCTTTGTCCAAA<br>A |
| PACRG-B-NG tagging | Forward | | ACTTGCTCGAACAATACGGTGGGGA<br>GGACGCGTACATCAACATTAAATACA<br>TGGTCCCGACGTATGAGTCATGCCTC<br>ATGCTTGTCTCGAAAGGTGAGGA |
| PACRG-B-NG tagging | Reverse | | CTGTGTATCTCTTTCTCTCCTTGTCTA<br>CTTTTATACACTCCTTGCTTTATCAGC<br>TTAACAGCAACGGAACGACGAGACTA<br>CCAATTTGAGAGACCTGTGC |
| PACRG-B KD | Forward | XbaI | GCtctagaTCCCGCCTACTGAGTTCA<br>GA |
| PACRG-B KD | Reverse | HindIII | CCCaagcttTGTTGATGTACGCGTCC<br>TCC |
